# Biocompatibility characterisation of CMOS-based Lab-on-Chip electrochemical sensors for in vitro cancer cell culture applications

**DOI:** 10.1101/2023.11.23.568427

**Authors:** Melina Beykou, Vicky Bousgouni, Nicolas Moser, Pantelis Georgiou, Chris Bakal

## Abstract

Lab-on-Chip electrochemical sensors, such as Ion-Sensitive Field-Effect Transistors (ISFETs), are being developed for use in point-of-care diagnostics, such as pH detection of tumour microenvironments, due to their integration with standard Complementary Metal Oxide Semiconductor technology. With this approach, the passivation of the CMOS process is used as a sensing layer to minimise post-processing, and Silicon Nitride (Si_3_N_4_) is the most common material at the microchip surface. ISFETs have the potential to be used for cell-based assays however, there is a poor understanding of the biocompatibility of microchip surfaces. Here, we quantitatively evaluated cell adhesion, morphogenesis, proliferation and mechano-responsiveness of both normal and cancer cells cultured on a Si_3_N_4_, sensor surface. We demonstrate that both normal and cancer cell adhesion decreased on Si_3_N_4_. Activation of the mechano-responsive transcription regulators, YAP/TAZ, are significantly decreased in cancer cells on Si_3_N_4_ in comparison to standard cell culture plastic, whilst proliferation marker, Ki67, expression markedly increased. Non-tumorigenic cells on chip showed less sensitivity to culture on Si_3_N_4_ than cancer cells. Treatment with extracellular matrix components increased cell adhesion in normal and cancer cell cultures, surpassing the adhesiveness of plastic alone. Moreover, poly-l-ornithine and laminin treatment restored YAP/TAZ levels in both non-tumorigenic and cancer cells to levels comparable to those observed on plastic. Thus, engineering the electrochemical sensor surface with treatments will provide a more physiologically relevant environment for future cell-based assay development on chip.

## Introduction

Advances in integrated circuit (IC) and microfluidic design have given rise to the miniaturisation of standard, laboratory-based tests on silicon microchips, in the form of Point-of-Care (PoC) diagnostics.^1–3^ Portable electronic devices allow rapid and sensitive data collection for diagnosis but also benefit from scalable and cost-effective manufacturing.^3^ A silicon-based sensor, which has gained significant popularity since its introduction by Bergveld in 1972, is the Ion-Sensitive Field-Effect Transistor (ISFET).^4,5^ Inherent pH sensitivity has driven its development as a biosensor for diagnostic tests including DNA, RNA, protein detection and cell-based assays on chip.^6,7^ But despite the development of ISFETs for cell-based assays with mammalian cells by several groups, the biophysical interactions and mechano-response of cells at the gate surface remains unappreciated as the focus of many studies remains on system design and applications.^8–11^

The ISFET was developed as an adaptation to the earlier MOSFET (Metal-Oxide Semi-conductor Field-Effect Transistor) where the date is tied to a reference electrode in contact with the aqueous solution and the gate oxide is replaced by an insulator.^4,5,12^ In unmodified CMOS technology, the gate is tied to the top metal of the process and the passivation layer is used as sensing membrane. The first ISFET design featured a silicon oxide passivation layer whose sensitivity to hydrogen ions rendered the ISFET a pH sensor.^4,5,12^ The reaction of silanol groups (Si-OH) at the gate interface determines the availability of H^+^ ions in solution and hence the resulting pH.^13^ Subsequently, implementation of ISFETs on standard Complementary Metal-Oxide Semiconductor (CMOS) technology and Moore’s law, has made it an inexpensive and scalable sensor for commercial production.^2,4,12,14,15^ The standard top material for CMOS passivation is Silicon Nitride (Si_3_N_4_). With unmodified CMOS integration, the Si_3_N_4_ passivation layer is retained making production rapid and low-cost by alleviating the need for extensive post-processing.

The first measurements on ISFETs, demonstrated by Bergveld, were based on ionic fluctuations and electrophysiological recordings of neurons. ^4,5,16^ A number of studies have developed ISFET sensors for cell-based assays, however, there is little focus on whether the interaction with a silicon-based interface has an effect on cell phenotype. Lehmann et al (2000), first reported a modified CMOS ISFET array with an Al_2_O_3_ passivation layer, where extracellular acidity was recorded in adenocarcinoma cells with a near-Nernstian sensitivity of 56mV/pH.^8^ Milgrew et al (2008), also reported a 16×16 ISFET array with a Si_3_N_4_ interface for detection of pH in fibroblast cultures.^9^ Finally, another high k-dielectric gate material, Ta_2_O_5_, has also been investigated with success in increasing sensor sensitivity.^10,11,17^ The variable materials which can be found at the interface of ISFET sensors have all been successfully implemented and demonstrate sensor sensitivity. However, the biophysical interactions at the Si_3_N_4_ interface remain unappreciated.

Bio-material design has been shown to play a central role in cell behaviour.^18,19^ Varying degrees of stiffness, roughness, patterning and elasticity contribute to cell adhesion and shape. Key regulators of mechanotransduction in mammalian cells are the transcriptional co-activators YAP/TAZ. YAP/TAZ translate biophysical changes in the microenvironment to transcriptional changes, allowing cells to be mechanically responsive to their surroundings. The activation of YAP/TAZ is most notably affected by mechanical microenvironmental changes, such as substrate stiffness, and the resulting formation of focal adhesions and cell shape.^20–22^ When activated, YAP/TAZ translocates from the cytoplasm to the nucleus where it can impact intracellular signaling to activate downstream signaling pathways, such as the Hippo pathway and Wnt signaling cascade.^22–24^ In particular, the relevance of its activity can be appreciated through its wider effects on cell proliferation, growth and metabolism.^22–24^

This study focuses on the biocompatibility of the passivation material, Si_3_N_4_, as a case study for CMOS-based electrochemical sensors. We show that non-tumorigenic and tumorigenic mammalian cells grown on ISFET arrays with a Si_3_N_4_ interface demonstrate lower attachment efficiency to Si_3_N_4_, when compared to attachment on standard tissue culture plastic. Higher YAP/TAZ concentration in the cytoplasm demonstrates lower activation on chip than on plastic. Expression of proliferation marker, Ki67, shows an increase on tumorigenic cells grown on Si_3_N_4_. Meanwhile, non-tumorigenic, epithelial cells cultured on Si_3_N_4_ ISFETs showed a similar profile of cell adhesion and proliferation as tumorigenic cells, however a slight increase in YAP/TAZ activation on chip was recorded. Surface treatment with Poly-L-ornithine and laminin (PLOL) rescued the proliferation and mechanotransduction profile of non-tumorigenic and tumorigenic cells to mimic the cell phenotype observed on cell culture plastic to a greater extent than other treatments investigated. YAP/TAZ and proliferation marker expression on collagen and fibronectin (FN)-treated chips did not improve the resemblance of the cell phenotype to the one observed on plastic in both normal and cancer cell cultures.

## Results and discussion

The lab-on-chip diagnostic device used in this study has been previously described in Moser et al (2018).^25^ The microchip cartridge harbouring a 4,368 ISFET pixel array is attached using a 3D printed clip mechanism onto a Printed Circuit Board (PCB) to integrate user functionality and digitization of output results. The PCB itself enables Bluetooth connection to an Android device with an application which allows inspection and storage of real-time ISFET output. The array holds a volume of 20 *µ*l in a clear, acrylic manifold and is connected to an Ag/AgCl reference electrode. The ISFET array used in this study, which is manufactured in unmodified CMOS technology with a Si_3_N_4_ passivation layer, limiting both production costs and signal decoupling from the sensor.^25^

### ISFET arrays versus tissue culture plastic for cell culture

#### Cell adhesion and morphogenesis on ISFET arrays

The experimental workflow used in this study is outlined in Figure 1a. Cells are cultured on plastic or ISFET chips for 48hours and subsequently immunofluorescently stained with markers for proliferation (Ki67), mechanotransduction (YAP/TAZ) as well as nuclear (Hoescht) and cytoskeletal stains (*α*-Tubulin, F-actin). Images obtained are parsed through an automated feature extraction pipeline is used to obtain quantitative values for cell shape, proliferation and mehanotransduction. Finally, we train a linear classifier to extract distinct cell shapes to assess heterogeneity in the cell population (Figure 1a).

**Figure 1:**
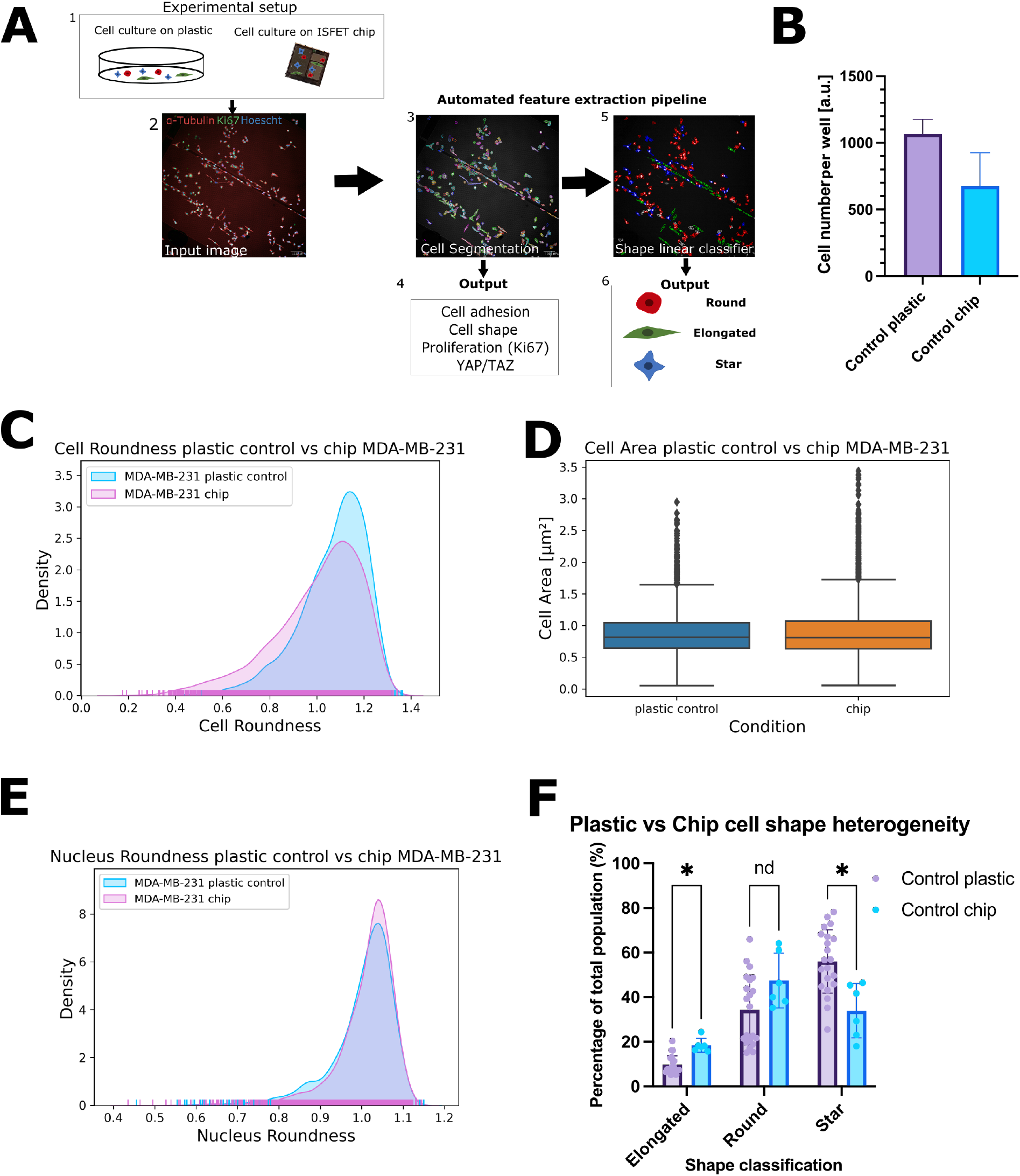
Experimental workflow and cell morphology of triple-negative breast cancer cell line, MDA-MB-231, on tissue culture plastic versus the Si_3_N_4_ ISFET gate surface. **(a)** Experimental setup of cell culture on plastic or ISFET chips followed by automated feature extraction pipeline from high-throughput confocal microscopy, using the Perkin Elmer Columbus software. A fluorescent confocal image is used as input for segmentation of nucleus and cell shape and extraction of shape, intensity, and texture feature. Selection of training data and extracted features allow classification of three different shape classes (round, elongated, star). **(b)** Number of cells attached per well, after 48 hours cell culture. N=6 chips, N=21 wells. **(c)** Kernel density estimation and corresponding rug plot of cell roundness for cells cultured on plastic 96-well cell culture plates and the ISFET array. **(d)** Cell area (*µ*m^2^) in cells cultured on plastic culture plates as a control versus the ISFET chip. **(e)** Kernel density estimation plot and corresponding rug plot of nucleus roundness for cells cultured on plastic culture plates or the ISFET chip. N = 4,062 single cells for all conditions. **(f)** Percentage of total population attached on plastic or Si_3_N_4_ ISFET arrays classified as elongated, round or star. Multiple t-test discovery indicated by asterisk (*) N=6 chips N=21 plastic wells. Statistics performed in python of GraphPad Prism version 9.5.0.

Firstly, we investigated if the ISFET array with a Si_3_N_4_ passivation layer affected cell adhesion or cell morphogenesis. We compared cell adhesion and cell shape on ISFET arrays manufactured in standard CMOS technology to cells cultured for 48 hours on commercially-available, 96-well culture plates. After 48 hours, we observed approximately 1000 cells/well attached to standard plastic cell culture vessels, reflecting 100% of initially plated cell numbers, as opposed to almost 50% on untreated, control chips (Figure 1b). Unpaired 2-tailed t-test statistics show that cell attachment on a Si_3_N_4_ passivation layer is significantly lower than on plastic microplates (p-value *<* 0.0001, difference between means -389.571 *±* 68.69) (Figure 1b).

To determine if cell shape or cytoskeletal organization is different in cells cultured on an ISFET array to those on standard tissue culture (TC), we quantified features describing morphology. The roundness of cells grown on plastic is significantly different to those grown on chip as the kernel density estimation (KDE) plot shows a higher peak at 0.14 a.u, which is also confirmed by 2-sample Kolmogorov-Smirnov (KS) statistical testing (p = 3.77e^-35^) (Figure 1c). The KDE plot kurtosis also indicates a greater variation in the distribution of cell roundness on chip. On the other hand, the area of cells grown on chip versus on plastic was not found to be significantly different, suggesting that solely the shape of cells is affected by the Si_3_N_4_ surface (Figure 1d). Nucleus roundness was significantly different on chip, with a greater number of single cells with a lower nucleus roundness as indicated by the spread of the KDE plot in Figure 1e. It is noteworthy, that we can distinguish the singular pixels of the ISFET array as rectangles. By examination of the *α*-Tubulin or F-actin, the direction of growth is seen to run parallel to the pixel axes, resulting in an elongation of cell shape (Figure 1a & Figure 1f). This is reflected in Figure 1f where the percentage of the total population classified as an elongated cell shape by a linear classifier is shown to be increased on ISFET chips as compared to plastic. Unpaired multiple t-test, confirms a discovery of differences in the percentage of total cells classified as elongated and star shape on chip which is likely to be attributed to the ISFET pixel architecture (Figure 1f). These results indicate that whilst there are subtle changes in cell roundness which may be related to the surface configuration of the ISFET interface having an impact on the cell shape, cell size remains the same on chip as on plastic.

#### Proliferation on ISFET arrays

To evaluate the effect of Si_3_N_4_ as a substrate for cell growth, we quantified the levels of proliferation marker, Ki67, in single cells^26^[41]. Figure 2a is representative of the localisation and intensity of Ki67 foci found in the nuclei of MDA-MB-231 cells on chip. Mean normalised Ki67/Hoescht intensity ratio of cells grown on plastic is 0.878 a.u. The ratio is 37% higher on chip. The cumulative distribution function (CDF) of normalized Ki67 intensity in cells grown on chip shows the point where the maximum difference can be observed (ks statistic = 0.244, p-value = 1.76e^-106^), indicating that there is a significantly different expression of Ki67 on chip as compared to cells on plastic (Figure 2b). Further inspection of the KDE panel reveals that a greater proportion of the population grown on chip are actively in any stage of the cell cycle (G1,G2,M) as indicated by the presence of Ki67 foci (Figure 2a & Figure2b). This confirms that there is no evident initiation of quiescence on chip, which could otherwise be distinguished through decreasing Ki67 levels.^26–28^

**Figure 2:**
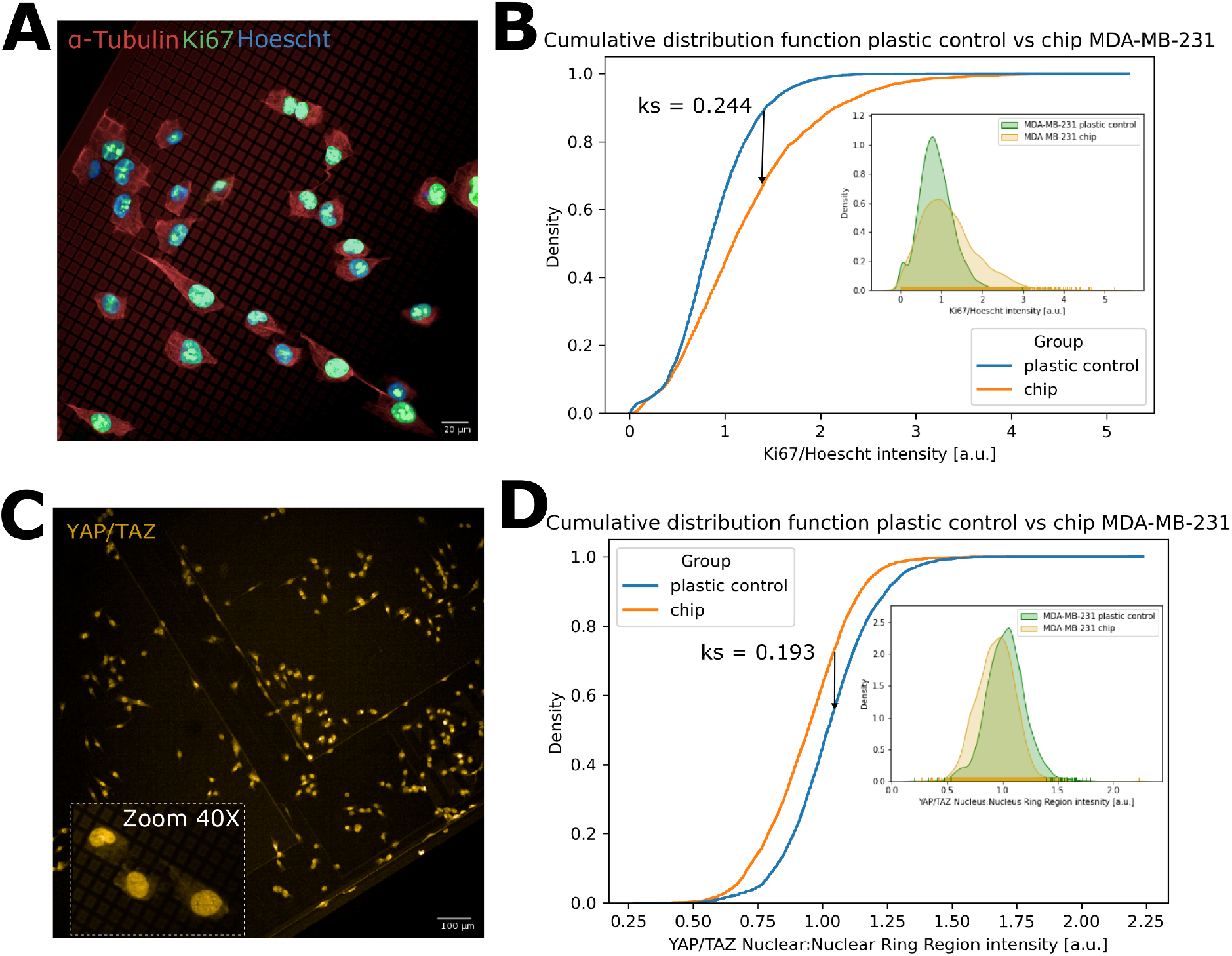
Characterisation of changes in proliferation and YAP/TAZ expression of Triple negative breast cancer cells, MDA-MB-231, on plastic cell culture plates versus the Si_3_N_4_ ISFET gate surface. **(a)** MDA-MB-231 cells cultured on the ISFET array with fluorescent antibody staining for alpha Tubulin (red), proliferation marker Ki67 (green) and Hoescht nuclear dye (blue). **(b)** Cumulative distribution function of Ki67 intensity, normalised to Hoescht dye intensity, for single cells cultured on plastic culture plates as a control versus ISFET chips. Panel of kernel density estimation and corresponding rug plot. **(c)** MDA-MB-231 cells cultured on the ISFET array and fluorescently stained for YAP/TAZ expression (yellow) with a zoom panel at 40X magnification. **(d)** Cumulative distribution function of YAP/TAZ intensity ratio in the nucleus versus the nuclear ring region in MDA-MB-231 cells cultured on plastic versus ISFET arrays. Panel of kernel density estimation and corresponding rug plot. Plastic control N = 21 wells or ISFET chip N = 6. N = 4,062 single cells for all conditions. All images obtained using the Opera Phenix (Perkin Elmer) and segmented with Columbus (Perkin Elmer). Kolmogorov-Smirnov statistics (Ks) performed using the scipy.stats module in Python.

#### Mechanotransduction on ISFET arrays

Most cells are responsive to mechanical forces which are a function of how cells interact with their microenvironment.^23^ We have previously observed that other materials, such as Si Nanoneedles, can significantly alter the dynamics of mechanically sensitive signalling pathways such as those regulating the YAP/TAZ mechanosensors.^20^ To evaluate whether culturing of breast cancer cells on chip also affected mechanically sensitive signalling pathways, we quantified nuclear localization of YAP/TAZ on cells cultured on ISFET arrays. When activated by mechanical stress, YAP/TAZ is translocated from the cytoplasm where it lies inactive, into the nucleus.^29^ By segmentation of the nucleus and the cytoplasmic area surrounding the nucleus, herein referred to as nuclear ring region, we calculated a YAP/TAZ ratio. Figure 2c is a representative, confocal image of cells on chip showing intensity of YAP/TAZ in the nucleus and cytoplasm. The CDF plot of YAP/TAZ intensity shows a significant difference between the two populations (p-value = 1.51e^-66^), which is also validated by the KDE panel where the peak of the population on chip is positively skewed (Figure 2d). Thus, YAP/TAZ distribution is on average higher in the nuclear ring region than in the nucleus in the chip population versus plastic resulting in a greater proportion of YAP/TAZ remaining inactive in the cytoplasm of cells grown on chip. As a result, cells cultured on chip are likely less mechanically responsive to the Si_3_N_4_ surface than they are on plastic.

### Extracellular matrix component (ECM) deposition on ISFET arrays

#### Cell adhesion and morphogenesis on ECM-treated ISFET arrays

In order to examine whether the biocompatibility of chips could be improved, chips were coated with rat tail collagen Type I, human FN or PLOL.^30–32^ Figure 3a depicts representative confocal images of chips with the relevant extracellular matrix deposits where cells are stained for cytoskletal marker, *α*-Tubulin, Ki67, YAP/TAZ and nuclear stain Hoescht. Firstly, a comparison of the number of cells on untreated plastic, untreated control chips and ECM-treated chips shows that cell adhesion and growth increased on all treated chips in comparison to control chips. Approximately double and triple cell numbers were observed on FN and collagen or PLOL surfaces, respectively, when compared to control chips (Figure 3b). In addition, collagen and FN treatment of chips resulted in rescue of cell adhesion to similar cell numbers adhered to standard cell culture plastic, whilst cell numbers on PLOL surpassed standard plastic vessels attachment efficiency (Figure 3b). However, it is noteworthy, that the number of cells attached to collagen and FN chips varied on a per chip level (Figure 3b).

**Figure 3:**
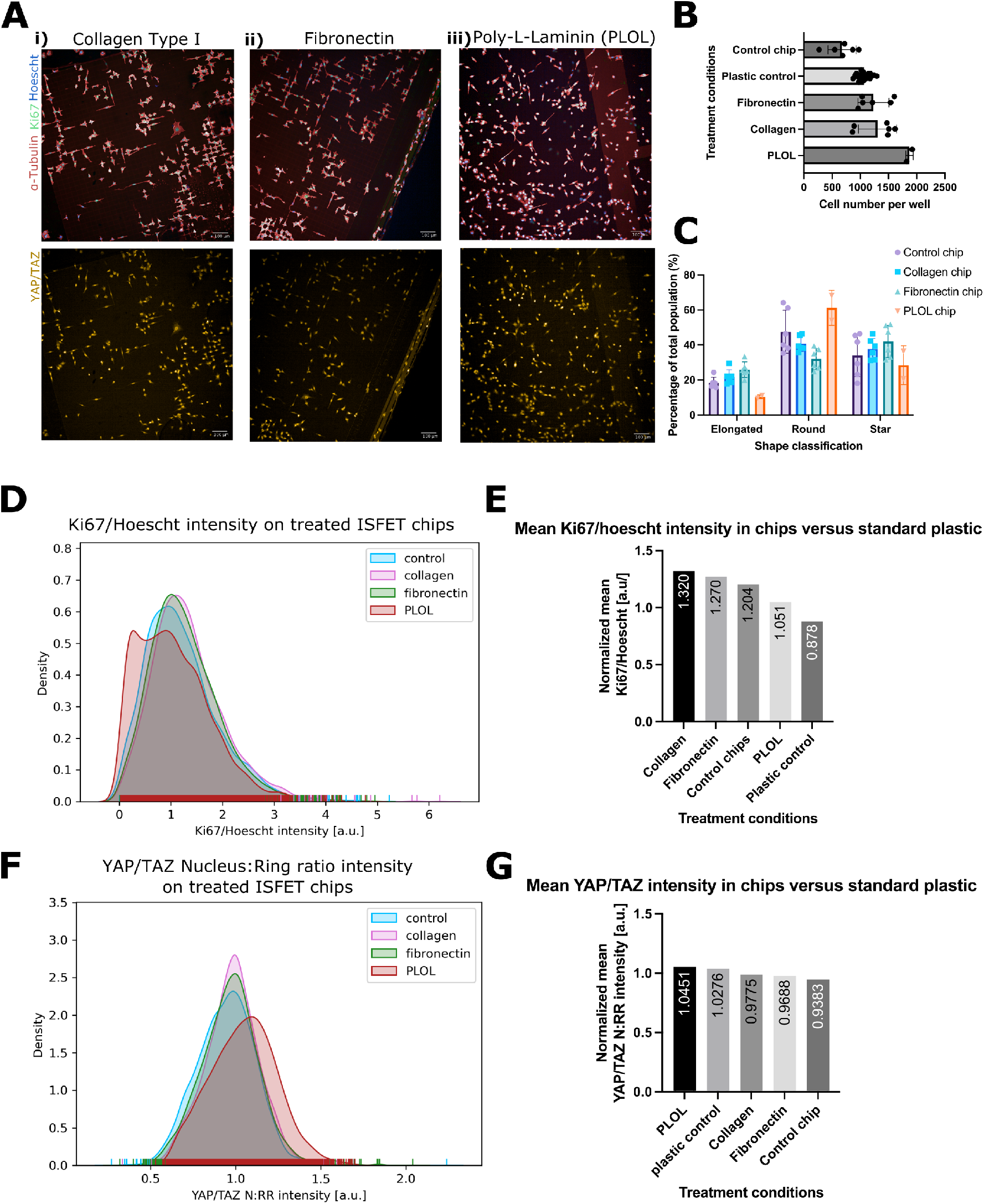
Cell adhesion, cell shape, proliferation and YAP/TAZ expression of cancer cells on Si_3_N_4_ interface treated with extracellular matrix components. **(a)** MDA-MB-231 attached to ISFET arrays with deposited **(i)** Collagen Type I (50 *µ*g/ml), **(ii)** Fibronectin (1*µ*g/ml) and **(iii)** PLOL (8 *µ*g/ml) deposited on ISFET surface prior to cell culture (from left to right). ISFETs are fixed and stained for cytoskeletal marker, *α*-Tubulin (red), proliferation marker, Ki67 (green), Hoescht nuclear dye (blue) in the upper panels and YAP/TAZ (yellow) in the lower panels. All images obtained on the Opera Phenix (Perkin Elmer). **(b)** Number of cells attached per well, after 48 hours, on all treated ISFET chips or untreated control chips versus standard culture plastic vessels. **(c)** Percentage of total population of cells attached on the respective ISFET chips whose shape were classified as elongated, star or round. **(d)** Kernel density estimation and corresponding rug plot for proliferation marker, Ki67, expression normalised to Hoescht dye intensity in cells cultured on control chips and treated chips. **(e)** Normalized mean Ki67/Hoescht intensity in treated chips or untreated chips versus standard culture plastic. **(f)** Kernel density estimation and corresponding rug plot for YAP/TAZ nuclear: nuclear ring ratio intensity on control chips or treated chips. **(g)** Normalized mean YAP/TAZ N:RR in treated chips or untreated chips versus standard culture plastic. N = 3,743 single cells for all KDE plots. All images obtained using the Opera Phenix (Perkin Elmer) and segmented with Columbus (Perkin Elmer). Kolmogorov-Smirnov statistics (Ks) performed using the scipy.stats module in Python.

Cytoplasm, nucleus shape and texture features were used to train a linear classifier to extract 3 distinct cell shapes in the MDA-MB-231 population: elongated, round and star. Strikingly, a higher percentage of the population of cells grown on PLOL are classified as round, with a corresponding decrease in the proportion of elongated cells, in comparison to control chips (Figure 3c). However, multiple t-test shows that there is no significant difference between the cell shape distributions of all chip configurations in comparison to untreated, control chips.

#### Proliferation on ECM-treated ISFET arrays

Figure 3d shows a KDE and corresponding rug plot of Ki67 intensity in chip configurations. A bimodal distribution can be distinguished in the PLOL-treated populations whilst all other configurations follow a positively skewed, unimodal distribution. Additionally, 2-sample KS test indicates that both collagen (p-value = 3.42e^-10^) and FN (p-value = 9.7e^-08^) groups show significantly different Ki67 expression as compared to the control (Figure 3d). When compared to standard cell culture plastic, PLOL-treated chips more closely resembled the proliferation profile observed under standard cell culture conditions with a normalized mean Ki67/Hoescht intensity of 1.051 a.u. (Figure 3e).

#### Mechanotransduction on ECM-treated ISFET arrays

To determine if the mechanical sensitivity of cells was impacted due to ECM treatment of ISFET chips, we quantified YAP/TAZ nuclear translocation in cells plated on chip with different substrates. Figure 3f shows all chip configurations following a similar trend of positive kurtosis as the control group, except the PLOL-treated population which is more normally distributed. Thus, the PLOL-treated group has a higher median YAP/TAZ intensity than all other configurations, indicating higher nuclear translocation of YAP/TAZ (Figure 3f). More specifically, the normalized mean YAP/TAZ N:RR intensity of PLOL-treated chips is found to be higher than recorded on plastic vessels indicating an increase in YAP/TAZ activation (Figure 3g). Collagen and FN-treated chips showed a moderately decreased YAP/TAZ activation to plastic (Figure 3g). However, statistical testing confirms that both the collagen and FN populations are also significantly different in YAP/TAZ intensity to the control chip group (2-sample KS, p-value = 8.04e^-16^, p-value = 6.10e^e-06^, respectively). Whilst all chip configurations improved efficiency of cell attachment without significantly skewing the cell shape heterogeneity of the MDA-MB-231 population, it can be seen that both proliferation and mechanoresponsiveness change dependant on the ECM-coating of chips when compared to an untreated Si_3_N_4_ interface. Most notably, PLOL-treated chips resulted in the highest number of proliferating cells, as indicated by Ki67 expression and a significant increase in the mechano-responsiveness of cells when compared to untreated Si_3_N_4_ chip surfaces.

### Focal adhesion formation on a Si_3_N_4_ interface

The formation of focal adhesions (FAs) is an essential component at the interface between cells and their extracellular microenvironment. They have a key role in anchoring cells to surfaces but also determine proliferation, signal transduction, as well as migration and are also upstream activators of YAP/TAZ through the Hippo pathway.^20,33^ We previously observed that Si-based nanoneedles inhibited the formation of FAs at contact sites.^20^ The impact of YAP/TAZ translocation in cells on chip lead us to investigate if FA morphogenesis was otherwise disrupted. FAs are found at the ends of stress fibres and can be identified through a scaffold protein, paxillin. Figure 4a, shows the expression of paxillin at FAs on mattek dishes (WT) and on ISFET chips (chips) treated with FN, obtained using Total Internal Fluorescence Microscopy (TIRF), to allow imaging of features closest to the coverslip. The left panels highlight the segmentation of FAs used to further characterise the pattern of FAs. The orientation of FAs under both conditions follows the same pattern, indicating that the patterning of the ISFET array, although affecting cell shape, does not affect the frequency distribution of FAs (Figure 4b). However, it can be distinguished that FAs formed on chip have a lower mean area, indicating stretching in their respective orientations (Figure 4c). This indicates that decreased activation of YAP/TAZ likely coincides with differences in FA morphogenesis on ISFET arrays.

**Figure 4:**
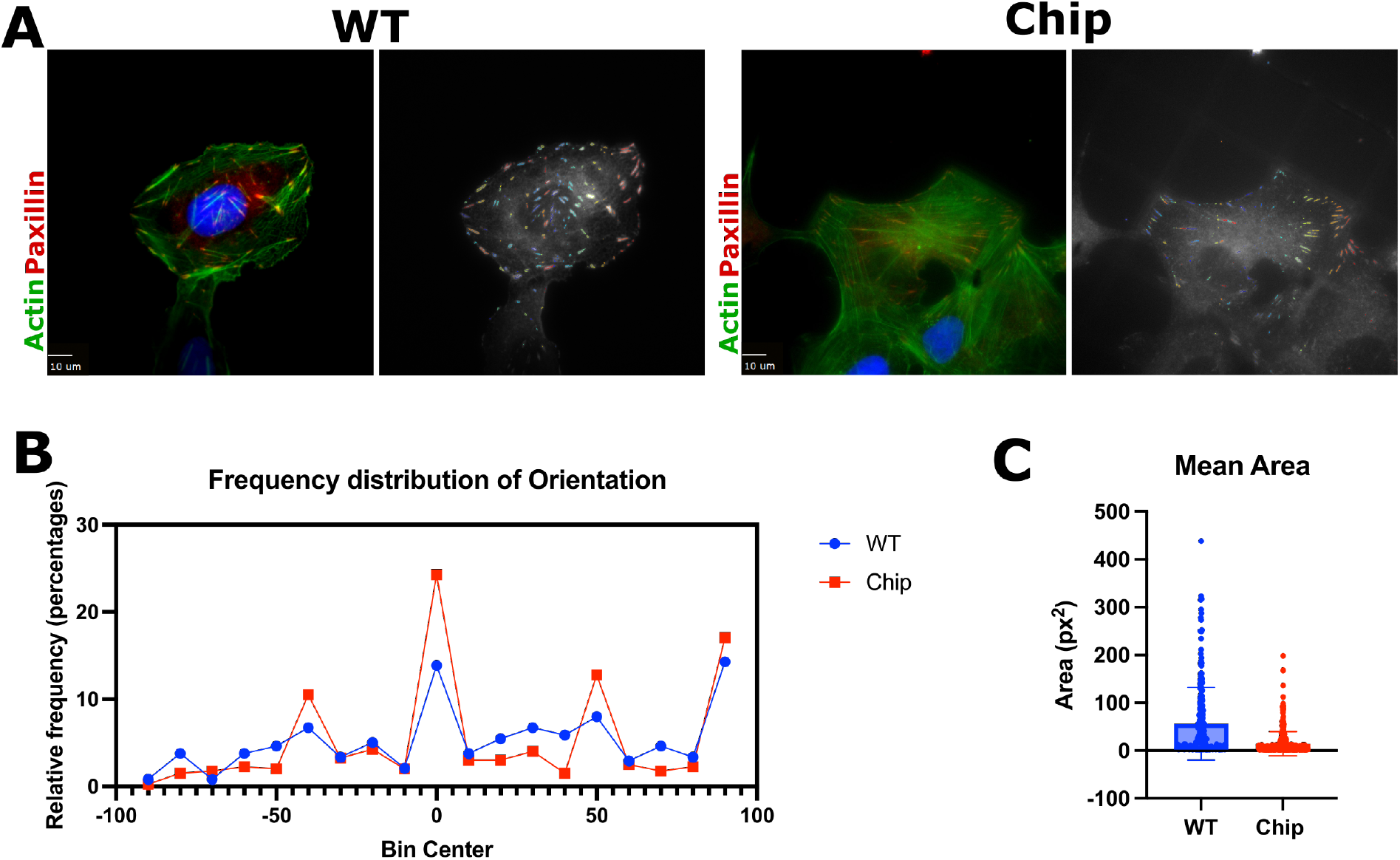
Focal adhesions of cancerous cells on standard tissue culture conditions (WT) and ISFET chip coated with fibronectin. **(a)** Cancerous cell line, U2OS CDK1 AS (osteosarcoma) was cultured on a standard tissue culture mattek dish (WT) or an ISFET array (Chip) both coated with fibronectin. In each panel: Right image shows paxillin stains of focal adhesions. Left image shows detection threshold segmentation of focal adhesions using the Focal Adhesion Analysis Server (FAAS). **(b)** Histogram of orientation of focal adhesions detection in WT versus chip conditions. **(c)** Mean area of single focal adhesions in WT versus on chip, as measured in square pixels. Images obtained on Total Internal Fluorescence (TIRF) microscope and quantiatively analysed using the Focal Adhesion Analysis Server. ^51^ Graphs and statistics performed with GraphPad Prism version 9.5.0.

### Cell morphogenesis, proliferation and mechanotransduction of non-tumorigenic cell culture on a Si_3_N_4_ surface of ISFET arrays

We also evaluated the biocompatibility of a non-tumorigenic cell line originating from mammary epithelial tissue, MCF10A. An assessment of the morphology of MCF10A cells on chip, indicates that whilst cells maintain their epithelial growth pattern, predominantly characterised by cell-cell contacts, when limited by the constraints of pixel architecture, they appear more elongated,than when cultured on standard culture vessels, whilst also maintaining cell contacts at either edge (Figure 5a). Quantitative evaluation of their elongation is represented in the ratio of cell width to length KDE plot where the greater number of cells concentrated at the tail of the distribution indicates the presence of more elongated cells on chip (Figure 5b). The significant difference between the two distributions is also confirmed with 2-sample KS statistics (p-value = 1.62e^-06^). Thus, the morphology of MCF10A appears to be affected by culturing on chip versus plastic as observed for cancer cells as well. The intensity of Ki67 was shown to be significantly different between plastic and chip in MCF10A cells (p-value = 1.12e^-16^), where a bimodal distribution in cells grown on plastic, versus an altered kurtosis in the chip population shows distinct differences in Ki67 expression of the two populations (Figure 5c). The distribution pattern of Ki67 on chip is comparable in both MDA-MB-231 and MCF10A cells, showing repeatability of results in different cell lines. Nonetheless, a greater maximum difference in Ki67 intensity occurs in MDA-MB-231 cells as evidenced by the ks statistic (Figures 2 & 5). Finally, YAP/TAZ intensity in the two populations shows that the peak of the chip distribution followed a negative skew and results in a significant difference in the two populations (p-value = 1.51e^-11^, ks statistic = 0. 077) (Figure 5d). In comparison to MDA-MB-231 cells, non-cancerous MCF10A cells YAP/TAZ intensity results show that the population distribution on plastic versus chip is similar in non-cancerous cells.

**Figure 5:**
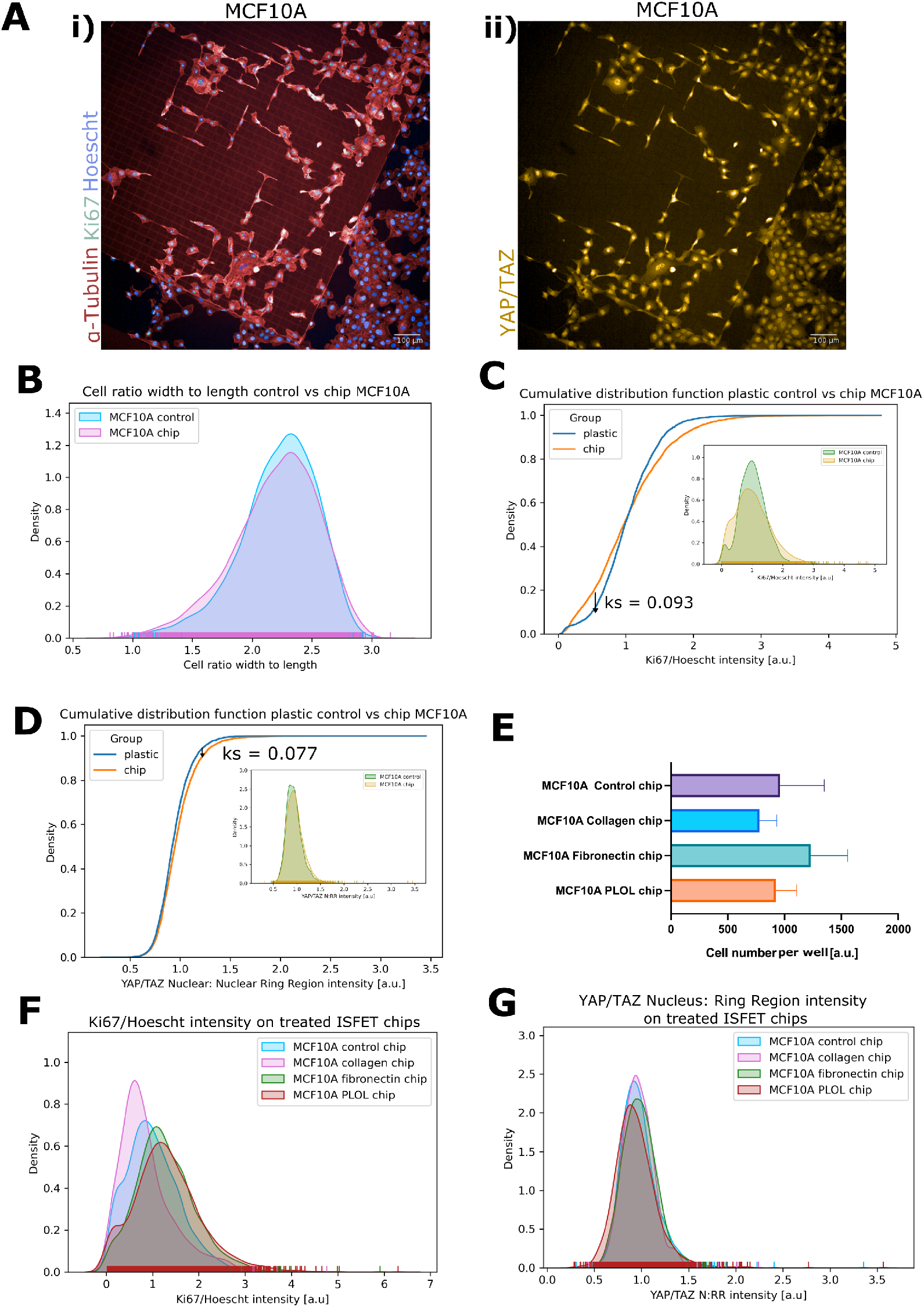
Non-tumorigenic mammary epithelial cells, MCF10A, morphology and behavioural response to culture on ISFET chips with Si3N4 interface. **(a)** MCF10A cells cultured on the Si_3_N_4_ interface of ISFET chips and fluorescently stained for **(i)** cytoskeletal marker *α*-Tubulin (red), proliferation marker Ki67 (green), Hoescht nuclear dye (blue) **(ii)** YAP/TAZ (yellow). **(b)** Kernel density estimation and corresponding rug plot of cell ratio width to length for MCF10A cells cultured on 96-well standard plastic cell culture plates as a control or ISFET chips. **(c)** Cumulative distribution function of Ki67 intensity, normalized to nuclear Hoescht dye intensity in cells cultured on plastic or ISFET chips. Panel of kernel density estimation plot showing single cell population density. **(d)** Cumulative distribution function of YAP/TAZ nuclear to nuclear ring ratio intensity in cells cultured on plastic or ISFET chips. Panel of kernel density estimation plot showing single cell population density. **(e)** Number of cells attached per well for ISFET chips with extracellular matrix deposition of Collagen Type I (50 *µ*g/ml), Fibronectin (10ng/ml) and PLOL (8 *µ*g/ml) or control chips with no additional deposition. (f) Kernel density estimation and corresponding rug plot for proliferation marker, Ki67, intensity normalised to nuclear Hoescht dye intensity in MCF10A cells cultured on control ISFET chips versus ISFET chip configurations. **(g)** Kernel density estimation and corresponding rug plot for YAP/TAZ nucleus to nuclear ring ratio intensity in MCF10A cells cultured on control, ISFET chips versus ISFET chip configurations. N = 4,282 single cells for all plastic and control ISFET chip experiments. N = 2,119 single cells for all chip configurations. All images obtained using the Opera Phenix (Perkin Elmer) and segmented with Columbus (Perkin Elmer). Kolmogorov-Smirnov statistics (Ks) performed using the scipy.stats module in Python.

### Cell morphogenesis, proliferation and mechanotransduction of non-tumorigenic cell culture on ECM-treated ISFET arrays

In order to increase the biocompatibility of the passivation layer, the same chip configurations as for cancerous cells (collagen, FN and PLOL) were deposited on chip, and attachment efficiency was evaluated. Figure 5e shows that whilst more cells on average attached onto the chips with a FN coating than on the untreated control chips, the difference is not significant, likely due to chip by chip variation. The KDE plot of Ki67 intensity for all chip configurations shows that the the MCF10A population of collagen-coated chips is positively skewed whilst FN and PLOL-treated are negatively skewed in comparison to the control group (Figure 5f). Collagen-coated chips show more actively proliferating cells, as indicated by the ratio of Ki67/Hoescht intensity, whilst FN-coated chips result in lower proliferation than the control group. All chip configurations have a significantly different proliferation profile to the control group. On the other hand, YAP/TAZ expression in all chip configurations shows that whilst there are differences in the peak location of the distributions which shows a different median, the overall shape of the distributions remains similar.

We conclude, that a similar behaviour can be observed on both native chips and ECM-treated chip configurations with both non-tumorigenic, mammary epithelial cells, MCF10A, and triple negative breast cancer cells, MDA-MB-231. Proliferation increased on chip for both cell types, whilst MDA-MB-231 cells also show an increase in YAP/TAZ intensity in the nuclear ring region which is not replicated in MCF10A cells where YAP/TAZ intensity slightly increases to indicate nuclear translocation. On the other hand, cell attachment efficiency increased in FN-treated chips for MCF10A cells, whilst attachment increased with all treatments as compared to native chips in MDA-MB-231 cells. This indicates that the choice of ECM deposition for cell-based assays is likely to be cell-type specific. PLOL treatment of chips restored both YAP/TAZ and Ki67 levels to be comparable to standard TC plastic in tumorigenic cells. Similarly, in non-tumorigenic MCF10A cells, PLOL rescued YAP/TAZ levels, however, untreated chips were most comparable to plastic in terms of proliferation levels. Both non-tumorigenic and tumorigenic cell lines showed a similar response in terms of proliferation and attachment on ISFET array chips.

The biocompatibility of Si_3_N_4_ has recently been scoped for biomedical applications, such as microspectroscopy and orthopaedic prosthetics, where studies have investigated cell growth and biochemical reactions at the cell-silicon interface^34–36^[19,20,21]. A range of cell types including, peripheral blood mononuclear cells (PBMNCs), osteosarcoma, osteoblastlike, stem cells, adipocytes, cardiomyocytes and fibroblasts have shown viability and growth of cells on silicon-based materials in discs or membrane-form.^34,35,37–42^ Notably, no DNA damage or reactive oxygen species response was reported with PBMNC culture, however, release of silicic acid and NH_4_^+^, affected cell metabolism in osteosarcoma^35,37^[20,22]. The release of ammonium ions could result in metabolism perturbations and proliferative capacity due to ammonia recycling through biosynthetic pathways which is known to occur in a breast cancer microenvironment.^43,44^ The release of ammonia was also shown in studies to be attributed to the chemical breakdown of Si_3_N_4_ when in contact with an aqueous solution, to form a layer of SiO_2_.^35,45^ Hence, it can be inferred that studies on Si_3_N_4_ can be reasonably extrapolated to a SiO_2_ interface. However, inflammatory cytokine, TNFa, was increased in culture with SiO_2_ but not Si_3_N_4_, likely resulting in distinct cell phenotypes on the two surfaces.^35^ Further evidence of prolonged growth of tumour xenografts in mice with Si_3_N_4_ implants showed the potential of the material as implantable sensors.^46^ Key to future applications is evidence of pH buffering at the interface has been reported in both mammalian and bacterial cultures which brings to question the cell-based assays which can be performed on Si_3_N_4_ surfaces.^37,41^

In line with these studies, we show that the Si_3_N_4_ gate surface of ISFET arrays, which can be successfully used as cell culture substrates, however, we observed significant differences in mechanotransduction of cells grown on ISFET arrays. The translocation of YAP/TAZ to the nucleus indicates activation, implying that the substrate stiffness of the ISFET surface is not similar to standard plastic culture vessels.^47^ Most importantly, this need not be seen as a limitation to the use of ISFET arrays, as the stiffness of standard culture vessels is well-beyond mammalian tissue norms.^47^ As a result, Si_3_N_4_ ISFET arrays are likely to represent a more physiological surface for the growth and survival of mammalian cells. Despite these observations, it has been noted that inconsistencies arise in study results due to variations in manufacturing and post-processing, such as thermal treatments, which alter the surface chemistry and can affect cell adhesion.^45,48,49^

Finally, we can significantly engineer the biocompatibility of the ISFET surface through the surface treatment with ECM components. We showed that all ECM treatments with collagen, FN and PLOL, resulted in a greater attachment efficiency in cancer cell cultures. Studies to mimic standard culturing conditions have investigated the use of PLOL and PLL(Poly-L-Lysine) coating on Si_3_N_4_ surfaces and also reported improved cell attachment.^32,50^ Notably, a lower growth rate and higher exit into quiescence was recorded for PC12 cells with neural properties.^50^ Here, we found that mean normalized Ki67/Hoescht ratio increases in breast cancer cells grown on chip as compared to plastic. This distinct difference may in fact arise due to faster proliferation in MDA-MB-231 cells on chip.^27^ With a higher concentration of Ki67 in single cells, we could speculate that faster proliferation may be linked to cell adhesiveness.^27^ In our study, PLOL treatment of ISFET arrays was shown to be overall most effective in increasing cell adhesion and rescuing mechanoresponsiveness of cells. Future applications of ISFET arrays for cell-based assays will require a consideration of the ISFET pixel architecture to optimise the surface for cell growth without introducing scaffolding which may affect both cell shape and mechanotransduction.

## Conclusion

This study investigated the effects of ISFET arrays with a Si_3_N_4_ passivation layer on cell adhesion, morphology, proliferation and mechanoresponsiveness of cancer cells and non-tumorigenic cells in vitro. The resulting cell phenotype indicates that the Si_3_N_4_ surface is a substrate on amenable to cell adhesion, however we observed a moderate increase in actively proliferating cells and significant differences in the mechanotransduction profile of tumorigenic cells. This can likely be attributed to a more physiological surface stiffness of ISFET arrays than standard plastic culture vessels. Overall, tumorigenic cells appeared to be more sensitive to the Si_3_N_4_ surface than non-tumorigenic cells. Introduction of ECM coating on ISFET surfaces, showed that PLOL treatment can modify the surface to restore mechanosensitivity to levels comparable to plastic in both tumorigenic and non-tumorigenic cells. As a result, differences in cell phenotype can largely be accounted for with the use of a PLOL coating on ISFET arrays. Based on our results we would propose that the Si_3_N_4_ gate interface of ISFET arrays are largely biocompatible.

**Table 1:** Summary of experimental results.

| Chip configuration/ Mean change versus TC plastic | Cell adhesion | Proliferation (Ki67) | Mechanotransduction (YAP/TAZ) |
| --- | --- | --- | --- |
| Cancer cells on chip | ↓ | ↑ | ↓ |
| Non-tumorigenic cells on chip | ↓ | ↑ | ↑ |
| Cancer cells on chip + Collagen | ↑ | ↑ | ↓ |
| Cancer cells on chip + FN | ↑ | ↑ | ↓ |
| Cancer cells on chip + PLOL | ↑ | ↑ | ↑ |
| Non-tumorigenic cells on chip + Collagen | ↓ | ↓ | ↑ |
| Non-tumorigenic cells on chip + FN | ↑ | ↑ | ↑ |
| Non-tumorigenic cells on chip + PLOL | ↓ | ↑ | no change |

## Experimental

### ISFET array configurations

The ISFET arrays used in these experiments were previously reported in Moser et al (2016). The ISFET arrays, also referred to as chips, are comprised of 4,368 pixels each. They are manufactured in standard 0.35*µ*m technology hence retaining the native silicon nitride, Si_3_N_4_ top passivation layer as the interface of the sensor with the electrolyte. ISFET arrays are planarised and cut by an external party to ensure surface roughness is minimal and uniform. Untreated, control chips are attached onto the well of a 96-well plate (PhenoPlate, PerkinElmer) using a small amount of lubricant and subsequently rinsed with PBS prior to use in cell culture.

#### Collagen treatment

Collagen Type I Rat Tail (3.4mg/ml, CORNING, Lot 0295002) was prepared to 50*µ*g/ml using 0.02M acetic acid (Fisher Scientific) for use as a surface treatment on chips. Chips were attached to the bottom of a 96-well plate, as for untreated chips, and incubated with 100*µ*l/well 50*µ*g/ml collagen solution, at 37 °C for 30mins within a 5% CO_2_ incubator. Subsequently, chips were washed twice with PBS and allowed to dry prior to cell culturing.

#### Fibronectin treatment

Fibronectin derived from human plasma (Sigma-Aldrich, Ref F0895) was diluted from a 0.1% solution to 1*µ*g/ml with PBS. Subsequently, chips in a 96 well-plate were incubated with 100*µ*l/well 1*µ*g/ml fibronectin solution at 37 °C for 30mins within a 5% CO_2_ incubator. Subsequently, chips were washed twice with PBS and allowed to dry prior to cell culturing.

#### Poly-L-ornithine and Laminin (PLOL) treatment

The protocol followed for PLOL-treatment of chips was previously reported by Hirata et al (2000). Chips in a 96-well plate were incubated with 250*µ*l Poly-l-ornithine 0.01% solution (100*µ*g/ml) (Sigma-Aldrich, Ref P4957) for 24 hours with UV irradiation in a sterile environment for 2 hours. Chips were washed twice with sterile, deionised water. Laminin solution from Engelbreth-Holm-Swarm murine sarcoma basement membrane (Merck/Sigma-Aldrich, stock 1mg/ml, Ref L2020) was diluted to a final concentration 8 *µ*g/ml and chips were incubated with 250*µ*l/well for 24hrs in a 37 °C, 5% CO_2_ incubator. Chips were washed twice at the end of treatment.

### Cell culture on plastic and chip

#### Maintenance & Passaging

Triple Negative Breast Cancer (TNBC) cell line, MDA-MB-231 was maintained in the following growth medium composition: Dulbecco’s Modified Eagle Medium (DMEM 1X) (4.5g/L D-Glucose, L-Glutamine and Pyruvate (Gibco), 10% heat-inactivated Fetal Bovine Serum (FBS) (Gibco) and 1% Penicillin/Streptomycin (Invitrogen). Mammary epithelial cell line, MCF10A, was maintained in the growth medium recipe in Table 2. All cell lines on both standard cell culture plates (Perkin Elmer) and ISFET array chips were maintained in 37°C, 5% CO_2_ incubators. Cells were maintained in T25 flasks and passaged when at 80% confluency. Existing cell medium was aspirated, each flask was washed with 5ml PBS and aspirated once more. For each flask, 1ml Trypsin-EDTA 0.25% (Gibco) was added and returned to the incubator until cells appeared detached. Once cells were detached, the volume was added to a falcon tube and centrifuged at 1000rpm for 5mins. The supernatant was aspirated and each pellet resuspended in an appropriate volume of the relevant growth medium. Cells were counted using a Cell Countess (Invitrogen) where 10*µ*l trypan blue and 10*µ*l of cell suspension were mixed and added to a cell counter slide. A solution of 4×10^3^ cells/ml was made for all cell lines. For conditions denoted “plastic control”, 250 *µ*l of the cell solution was added per well in a 96 well Phenoplate (Perkin Elmer). For all conditions on chip, 250 *µ*l of the cell solution was added to each well of a 96 well Phenoplate with the relevant chip affixed to the bottom surface. All conditions were cultured for 48hrs in a 37°C, 5% CO_2_ incubator.

**Table 2:**
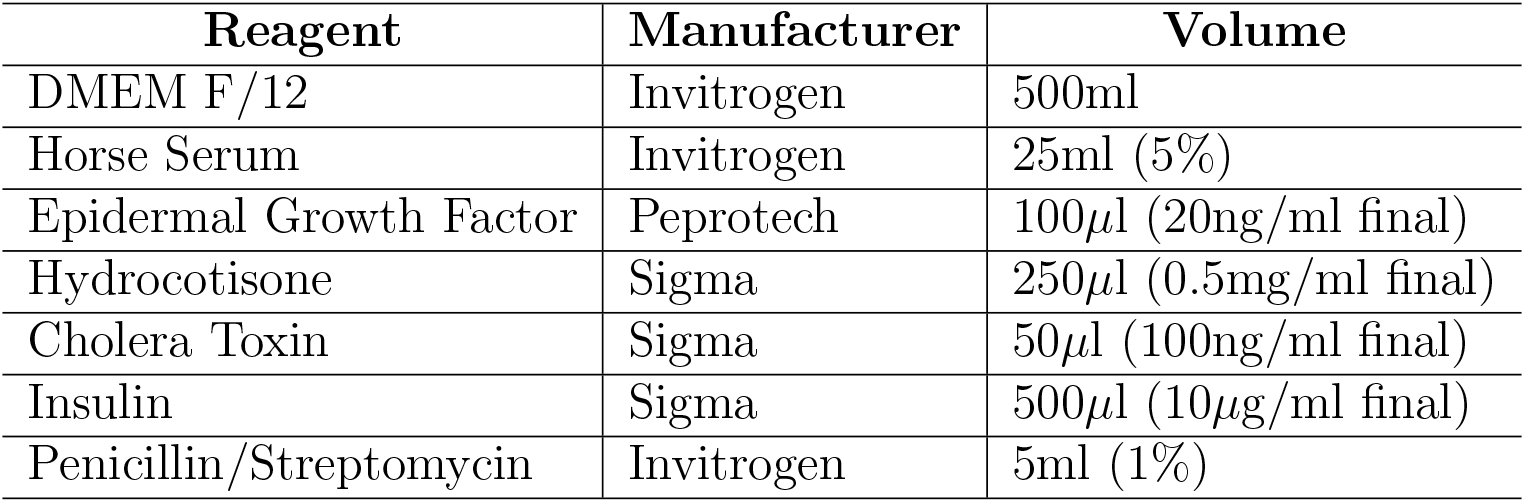
MCF10A growth media.

### Focal adhesions

Glass-bottom mattek dishes were coated with fibronectin derived from human plasma (Sigma-Aldrich) at a concentration of 10*µ*g/ml with incubation steps as described previously. ISFET chips were coated with 1*µ*g/ml as for all other chip experiments. Once treatment was completed, each mattek dish or ISFET chip was incubated with 20,000 cells/well overnight. Cells used in this experiment were U2OS CDK1AS-mCherry^52^ (a kind gift from Dr. Helfrid Hochegger, University of Sussex), an osteosarcoma cell line which were maintained in DMEM (Gibco) supplemented with 10% FBS (Gibco) AND 1% Penicillin/Streptomycin. Cells were maintained, passaged and counted as described in sub-section *Maintenance & Passaging*. Subsequently, experiments were fixed with 4% PFA, permeabilised and blocked as described in section *Immunofluorescence staining and confocal microscopy* with the use of anti-Paxillin (BD Biosciences, Cat 610052) as the primary antibody (Table 3). Focal adhesions were imaged using the TIRF (Total Internal Reflection Fluorescence) microscope. Images were processed and quantitative values extracted using the Focal Adhesion Analysis Server found at https://faas.bme.unc.edu/.^51^

**Table 3:** Primary and secondary antibodies for immunofluorescence staining.

| Primary Antibody | Manufacturer | Dilution | Secondary antibody | Manufacturer | Dilution |
| --- | --- | --- | --- | --- | --- |
| Rabbit Ki67 | abcam | 1:250 | Goat anti-rabbit IgG (H+L) Alexa Fluor 488 | Invitrogen | 1:1000 |
| Mouse YAP (63.7) | Santa Cruz technology | 1:1000 | Goat anti-mouse IgG (H+L) Alexa Fluor 568 | Invitrogen | 1:1000 |
| Rat α-Tubulin | BioRad | 1:1000 | Goat anti-rat IgG (H+L) Alexa Fluor 647 | Invitrogen | 1:1000 |
| Mouse Paxillin | BD Biosciences | 1:1000 | Goat anti-mouse IgG (H+L) Alexa Fluor 647 | Invitrogen | 1:1000 |

### Immunofluorescence staining and confocal microscopy

After 48 hrs, all conditions were fixed by removing the cell medium and adding a final concentration of 4% PFA for 15mins at room temperature. Wells were washed twice with PBS. Cells were permeabilised and blocked with 0.2% Triton-X for 12mins at room temperature, washed twice with PBS, followed by 2% Bovine Serum Albumin (BSA) for 1hr at room temperature. BSA solution was aspirated and primary antibodies indicated in Table 3 were added at the relevant dilutions in an antibody mix solution (0.5% BSA, 0.01% Triton-X and PBS), 100*µ*l/well were added and incubated overnight at 4 °C. The solution was then aspirated and wells washed twice with PBS before adding the relevant secondary antibodies in an antibody mix solution (100*µ*l/well) for 2hrs at room temperature (Table 3). Subsequently, wells were washed thrice with PBS. Sequential staining was performed where mouse and rat antibodies were utilized in the same wells. Hoescht dye and 647 Phalloidin-Atto 1:5,000 (where an *α*-Tubulin primary antibody was not used previously) was added in antibody mix solution for 10mins at room temperature. For focal adhesion work 488 Alexa Fluor Phalloidin 1:1,000 was used. Wells were washed once and left in PBS with NaN_3_. In order to image the chips, each chip was flipped to face downwards in fresh wells before imaging on the Opera Phenix high-throughput confocal microscope (Perkin Elmer).

### Image processing and segmentation

All image segmentation and feature extraction was performed using the Columbus software version 2.9.1 (Perkin Elmer). Confocal images obtained were taken in Z stacks and were maximally projected before segmentation. Hoescht staining of the nucleus was used to obtain nuclear roundness, with to length ratio, area and dye intensity. The same measurements were made for cytoplasmic and overall cell shape using Phalloidin dye or *α*-Tubulin for the cytoskeleton. Subsequently, intensity for proliferation marker, Ki67, was obtained for the nucleus area and normalized to the intensity of the Hoescht dye in the nucleus (Ki67/Hoescht ratio). YAP/TAZ intensity was measured in both the nucleus and a nuclear ring region surrounding the nucleus to create a ratio. A linear classifier was used to extract three shape populations (elongated, round and star shape) using all previously segmented features including roundness, area and SER texture as training data.

### Data analysis

Cell shape measurements of MDA-MB-231 cells were obtained from two independent experiment sets of 96 well plates and hence all values were normalized to the relevant feature’s plate average before further processing. The same was applied to experiments with MCF10A. Single cell values were used for all experiments. An equivalent number of single cells were sampled at random for all conditions. Data wrangling, plots and statistics of cell shape features, Ki67 and YAP/TAZ expression were performed using Python version 3.8.5 and packages scipy, matplotlib and seaborn. Number of cells attached per well or per chip, texture features and linear classifier data were plotted using GraphPad Prism version 9.5.0.

## Acknowledgement

The work of M.B., N.M., P.G., and C.B. in this project was supported by the CRUK Convergence Science Centre at The Institute of Cancer Research, London, and Imperial College London (A26234). The authors thank Filippos Maniatis, MSc student (Imperial College London), for his contribution in preliminary work towards the realisation of this project.

